# Modelling the role of LHCII-LHCII, PSII-LHCII and PSI-LHCII interactions in state transitions

**DOI:** 10.1101/2019.12.19.882886

**Authors:** W. H. J. Wood, M. P. Johnson

## Abstract

The light-dependent reactions of photosynthesis take place in the plant chloroplast thylakoid membrane, a complex three-dimensional structure divided into the stacked grana and unstacked stromal lamellae domains. Plants regulate the macro-organization of photosynthetic complexes within the thylakoid membrane to adapt to changing environmental conditions and avoid oxidative stress. One such mechanism is the state transition which regulates photosynthetic light harvesting and electron transfer. State transitions are driven by changes in the phosphorylation of light harvesting antenna complex II (LHCII), which cause a decrease in grana diameter and stacking, a decreased energetic connectivity between photosystem II (PSII) reaction centres and an increase in the relative LHCII antenna size of photosystem I (PSI) compared to PSII. Phosphorylation is believed to drive these changes by weakening the intra-membrane lateral PSII-LHCII and LHCII-LHCII interactions and the inter-membrane stacking interactions between these complexes, while simultaneously increasing the affinity of LHCII for PSI. We investigated the relative roles and contributions of these three types of interaction to state transitions using a lattice-based model of the thylakoid membrane based on existing structural data, developing a novel algorithm to simulate protein complex dynamics. Monte Carlo simulations revealed that state transitions are unlikely to lead to a large-scale migration of LHCII from the grana to the stromal lamellae. Instead, the increased light harvesting capacity of PSI is largely due to the more efficient recruitment of LHCII already residing in the stromal lamellae into PSI-LHCII supercomplexes upon its phosphorylation. Likewise, the increased light harvesting capacity of PSII upon dephosphorylation was found to be driven by a more efficient recruitment of LHCII already residing in the grana into functional PSII-LHCII clusters, primarily driven by lateral interactions.

**Statement of significance:** For photosynthesis to operate at maximum efficiency the activity of the light-driven chlorophyll-protein complexes, photosystems I and II (PSI and PSII) must be fine-tuned to environmental conditions. Plants achieve this balance through a regulatory mechanism known as the state transition, which modulates the relative light-harvesting antenna size and therefore excitation rate of each photosystem. State transitions are driven by changes in the extent of the phosphorylation of light harvesting complex II (LHCII), which modulate the interactions between PSI, PSII and LHCII. Here we developed a novel algorithm to simulate protein complex dynamics and then ran Monte Carlo simulations to understand how these interactions cooperate to affect the organization of the photosynthetic membrane and bring about state transitions.

## Introduction

The plant chloroplast thylakoid membrane is the site of the light reactions of photosynthesis. Here, the integral membrane protein complexes; light-harvesting complex II (LHCII), photosystem II (PSII), cytochrome *b*_*6*_*f*(cyt*b*_*6*_*f*), photosystem I (PSI) and ATP synthase carry out light harvesting and electron and proton transport, culminating in the synthesis of NADPH and ATP, which are utilized in the stroma for CO_2_ fixation. The thylakoid membrane is partitioned into the grana and stromal lamellae regions (1–3). The grana are cylindrical stacks of 2-30 appressed membranes 400-500 nm in diameter, enriched in LHCII and PSII. The stromal lamellae are flat sheet like structures that wrap helically around the grana interconnecting one stack to another and are enriched in PSI and ATP synthase (4–6). The cyt*b*_*6*_*f* complex is evenly distributed between the two domains (1, 7, 8). The division of the thylakoid into the grana and stromal lamellae depends on the presence of positively charged counterions to overcome the net negative charge on the membranes (9, 10). In the presence of cations attractive lateral interactions between PSII and LHCII drive the lateral segregation of these complexes from PSI and ATP synthase (10–12). In the presence of screening levels of cations attractive van der Waals and hydrophobic interactions between the flat stromal surfaces of these PSII-LHCII macrodomains then allow these complexes to self-associate with PSII-LHCII regions in neighbouring membranes to form 3-dimensional grana stacks (11, 13, 14). An entropic driving force through the influence of macromolecular crowding may also contribute to grana stacking (15). At the molecular level, the interaction of two LHCII trimers across the stromal gap has been calculated to be of the order 4 k_B_T where k_B_ is Boltzmann’s constant and T is the temperature in K (16).

The light environment of land plants fluctuates by orders of magnitude over time scales as little as a second. Such dramatic variations can cause mismatches between the amount of light absorbed and its utilisation in photosynthesis leading to metabolic and oxidative stress that lowers plant growth and productivity (17, 18). Fortunately plants have evolved a complex repertoire of mechanisms that can alter the efficiency of light harvesting and electron transfer in the thylakoid membrane (19). One such mechanism is the state transition which balances the relative excitation level of PSI and PSII to optimise photosynthesis (20, 21). The relative excitation balance between PSI and PSII varies with light spectral quality due to their differing absorption spectra and may also be adjusted to account for differences in the required ratio of linear (PSII and PSI) to cyclic (PSI only) photosynthetic electron transfer (22, 23). State transitions are regulated by the redox state of the photosynthetic electron transfer chain (24). When PSII is overexcited relative to PSI, the Q_p_ site on the cyt*b*_*6*_*f* complex is occupied by plastoquinol (PQH_2_) leading to activation of the STN7 kinase (25). STN7 phosphorylates the stromal-facing N-terminus of the LHCII subunits LHCB1 and 2, weakening its tendency to associate with PSII and increasing its affinity for PSI (State II) (26). In this way the relative light harvesting antenna sizes of PSI and PSII are altered to equalise their relative excitation rates restoring redox homeostasis of the electron transfer chain. When PSI is overexcited relative to PSII the PQ pool is oxidised and STN7 is inactivated and LHCII is dephosphorylated by the constitutively active phosphatase TAP38 (State I) (27, 28). Alternatively State I can also induced by simultaneous overreduction of the PQ pool and the chloroplast stroma in high light through the inhibitory effect of thioredoxin (and /or ΔpH) on STN7 (29, 30).

While the LHCII phosphorylation is clearly established as the mechanistic basis of state transitions it remains unclear precisely how the thylakoid macro-organization changes to bring about the relative change in the PSI/ PSII antenna size. STN7-dependent phosphorylation of LHCII causes a reduction in grana diameter and the number of membrane layers per stack while simultaneously increasing the number of grana per chloroplast (23, 31). Puthiyaveetil *et al*. (14) calculated, by solving the Poisson-Boltzmann equation for average surface charge across the grana, the changes in the stacking forces due to LHCII phosphorylation. They found that at physiologically relevant cation concentrations the balance between attractive van der Waals and hydrophobic interactions and repulsive electrostatic interactions is carefully poised. The net result is that increasing the negative surface charge density on LHCII by phosphorylation is sufficient to weaken the lateral and stacking interactions that sustain grana organization (13, 14). These changes bring about a reduction in grana dimeter and effectively increase the contact area between the grana and stromal lamellae, which may facilitate the exchange of LHCII between the domains (23, 31, 32). The changes in thylakoid stacking were also found to weaken the connectivity between PSII reaction centres, again contributing to the relative shift in antenna size (33). In addition to LHCII phosphorylation, molecular recognition through the presence of specific binding sites on PSI is also important (20, 34). Binding of Phospho-LHCII (P-LHCII) to PSI via the PsaL/ O and H subunits gives rise to the PSI-LHCII supercomplex for which a high resolution structural model has been solved (35, 36). A second probably weaker binding site for P-LHCII involving the LHCI (LHCA1-4) subunits of PSI also exists (37, 38). The *psal* and *lhca* mutants of *Arabidopsis* thus largely lack state transitions despite unchanged or even increased levels of LHCII phosphorylation (34, 37). Early structural analysis using freeze-fracture electron microscopy revealed that LHCII migrates from the grana to stromal lamellae upon transition to State II in isolated thylakoids, consistent with a decrease in the chlorophyll *a/b* ratio of this domain (39). However, later biochemical studies where state transitions were first induced in leaves prior to thylakoid isolation and those using *in situ* fluorescence microscopy suggested PSI and LHCII encountered one another in the margins where the two domains meet and exchange of LHCII between domains was very limited (40, 41). However, recent cryo-electron tomography data has cast doubt on the existence of a defined margins region with intermediate composition relative to the grana and stromal lamellae (8). Moreover, it is now clear that a population of LHCII energetically connected to PSI resides in the stromal lamellae even in State I and that the transition to State II rather increases the proportion (42). The relative importance of the changes in thylakoid membrane stacking and changes in the lateral interactions brought about by LHCII phosphorylation remain unclear and indeed they may fulfil distinct as well as complimentary roles in regulating electron a transfer and light harvesting

Given the unanswered questions that remain on the precise mechanism of state transitions new approaches are needed to clarify how the altered balance of forces upon LHCII phosphorylation affect the thylakoid macro-organization. The structural model of the thylakoid membrane at the individual protein complex level presented here was made possible by recent breakthroughs in elucidating thylakoid organization and dynamics using in atomic force microscopy (AFM) (7, 23, 43) and structured illumination microscopy (SIM) (31, 44), and publication of high-resolution structures of the key protein complexes involved in the light reactions (45–50). Recently developed algorithms, which made the calculation of whether two or more particles spatially overlap tractable, were also essential (see methods). Thanks to these advances, we were able to model the behaviour of protein complexes using structures obtained directly from the Protein Data Bank whereas previous studies relied on simplified protein geometries (16, 51). Previous models have revealed that relatively weak (2 K_B_T) interaction between LHCII trimers, including strong and moderately bound trimers of the PSII-LHCII supercomplex, give rise to complex emergent behaviour including the formation of crystalline domains of PSII and LHCII (51, 52). This model extends those of Schneider and Geissler., (16) and Lee et al., (51), to include cyt*b*_6_*f*, PSI and ATP synthase as well as introducing separate thylakoid regions: the grana and the stromal lamellae. We used this molecular model as a basis for Monte Carlo simulations to investigate the role of specific lateral intermolecular interactions and changes in grana stacking forces in modifying the distribution of photosynthetic complexes between the two domains and on the connectivity of LHCII with PSI and PSII. In particular, we address the question of whether the lateral and stacking interactions have functionally distinct or cooperative roles in state transitions. In order to achieve this, we developed a lattice-based algorithm for overlap detection of particles of arbitrary geometry an area that remains an active area of research. This algorithm was shown to be computational effective for a large number of protein complexes, rendering feasible the mesoscale Monte Carlo simulation protein structures directly from the Protein Data Bank.

## Materials and Methods

### Construction of the thylakoid model

A model of the thylakoid membrane was constructed using high-resolution structures of plant photosynthetic complexes, obtained from the Protein Data Bank (PDB accession codes: PSII-LHCII (C_2_S_2_M_2_): 5XNL (48), PSII-LHCII (C_2_M_2_): 3JCU (49), cyt*b*_6_*f*: 6RQF (45), LHCII: 1RWT (50), PSI-LHCI: 4Y28 (53), ATP synthase: 6FKF (46), on a square lattice with a spacing of 1 nm (Fig 1a). The densities of particles in the grana (dimeric PSII-LHCII and dimeric cyt*b*_*6*_*f*, 1.14 x 10^−3^ nm^−2^) and monomeric PSI-LHCI in the stromal lamellae (2.45 × 10^−3^ nm^−2^) were determined in a previous AFM study (23). We assumed a 1:1 ratio of PSII and PSI reaction centres (1) and a 1:1 ratio of C_2_S_2_ to C_2_S_2_M_2_ PSII-LHCII dimers (23). A number of free LHCII particles were added to the grana area to a total of 5 LHCII (including strong and moderately PSII-bound) per PSII reaction centre (54). cyt*b*_*6*_*f* particles were distributed between the grana and stromal lamellae at a ratio of at an equal area density to a total of 0.5 times the number of PSII reaction centres (1, 7). The ATP synthase was added to the stromal lamellae at a density of 0.7 of the number of PSII reaction centres (1, 55). Monomeric PSII and free LHCII were added to the stromal lamellae at a density of 10 % and 30 % of the total (PSII and LHCII respectively) (1). The radius of the stromal lamellae (*r*_*sl*_) was calculated using the equation

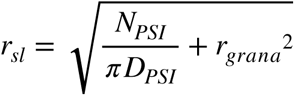

**Figure 1.**
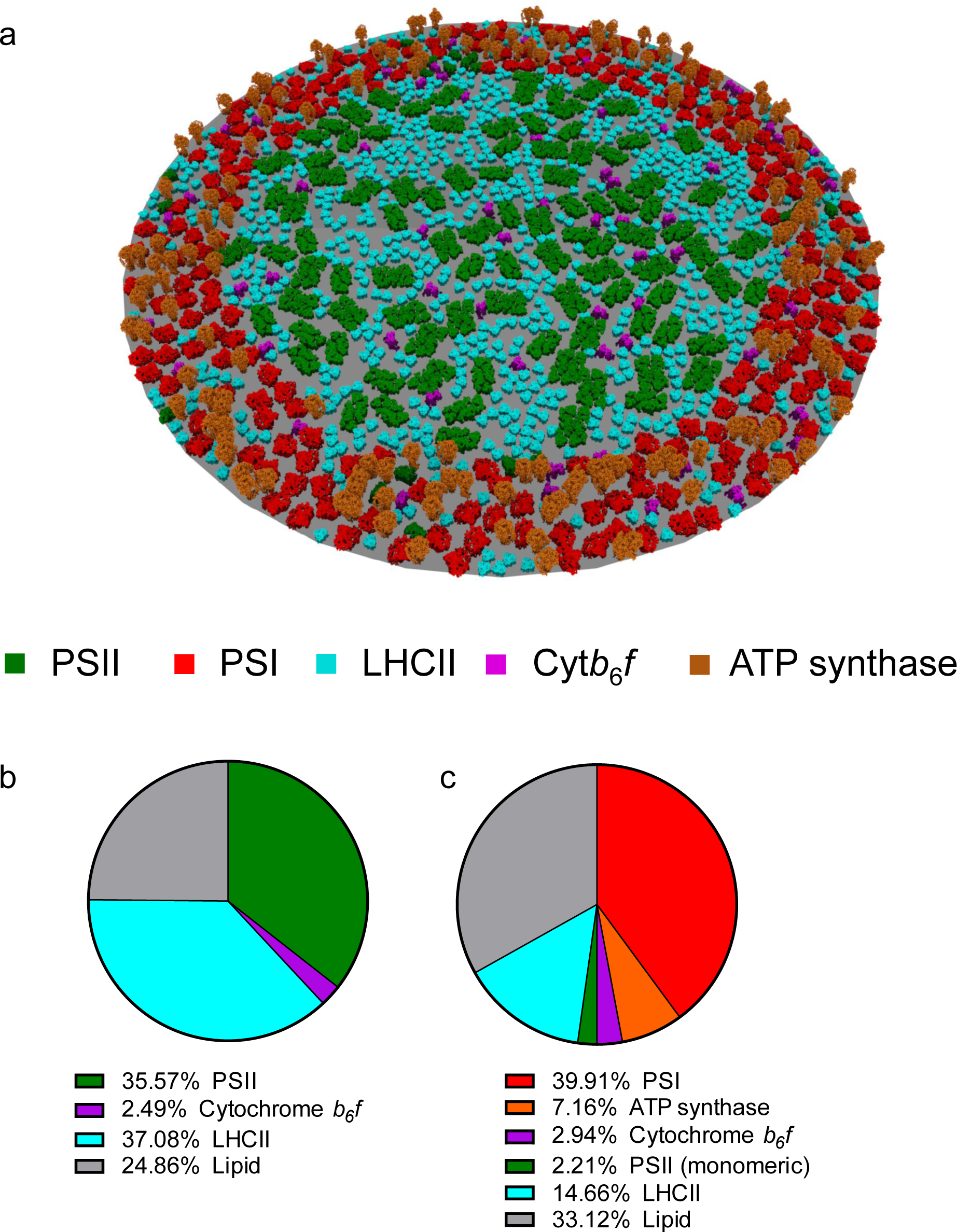
A structural model of the thylakoid membrane. a, A visual representation of the model. b, The percentage of grana area occupied by constituents of the grana. c, The percentage of stromal lamellae area occupied by constituents of the stromal lamellae.

Where r_grana_ is the grana radius, N_PSI_ is the number of PSI particles, and D_PSI_ is the density of PSI in the stromal lamellae.

### Monte Carlo simulations

LHCII lateral and stacking interactions were modelled using the potentials described in Schneider and Geissler (16) except that intra-layer interactions were nonspecific: we did not specify an angular difference threshold for binding. This is because specific interactions were considered from the outset by including C_2_S_2_M_2_ and C_2_S_2_ PSII-LHCII supercomplexes in the model in equal numbers. We note that the S and M LHCII trimers of C_2_S_2_M_2_ and C_2_S_2_ PSII-LHCII are not directly involved in state transitions *in vivo* (56). We introduced a PSI-LHCII interaction, based on Pan *et al*., (36), when LHCII was phosphorylated. This was modelled as a square well interaction and varied between 2 and 16 k_B_T with a distance threshold of 1 nm. Simulations included two grana layers of equal composition. Equilibrium properties of the thylakoid membrane system were sampled using the standard Metropolis-Hastings algorithm. For each perturbation in the Monte Carlo simulations, a particle was selected at random from one of the two layers, and moved by plus or minus 1 nm (lattice spacing) in x or y and rotated by plus or minus π/33. 10^7^ perturbations were made for each simulation and sampling was only carried out when the system was in equilibrium (Supplementary Fig. 1). Overlap of particles was forbidden during simulations. Spatial overlap was implied when the total number of lattice sites occupied by all particles was less than the sum of the number of lattice sites occupied by individual particles (Supplementary Fig. 2). A thorough discussion is given in Appendix 1. Test simulations were performed with only LHCII complexes (Supplementary Fig. 3) or with only LHCII and PSI complexes to test PSI-LHCII interactions.

### Diffusion in protein complex dynamics

Translational (D_T_) and rotational (D_R_) diffusion coefficients of protein complexes in the thylakoid membrane were calculated according to Saffman and Delbrück (57):

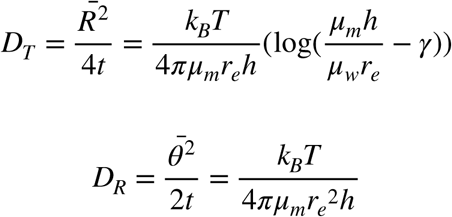

where *μ*_*m*_ = 200 *m Pa s* (based on the range of viscosities of lipid membranes of 160-260 mPa s(58)) and *μ*_*w*_ = 1 *m Pa s* are the viscosities of the lipid membrane and water respectively, *h* = 4 nm is the thickness of the lipid bilayer, *γ* ≈ 0.577 is the Euler-Mascheroni constant and *r*_*e*_ is the effective radius of the particle

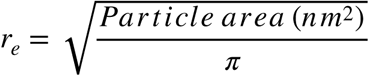

The effective radius therefore makes the simplifying assumption that the particle is circular. In diffusion simulations, translations were ± 1 nm (in x and y directions) and rotations were ± π/33 radians (π/33 radians was chosen because this results in movements to only neighbouring lattice site for larger particles). A single time step of 1 μs was used. To compensate for the variety of diffusion coefficients, we used a probabilistic approach whereby translations and rotations where accepted given the following criteria:

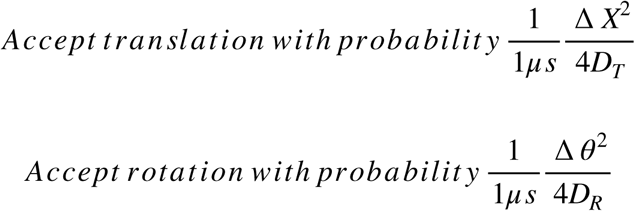

That is, movements were accepted with a probability defined by the predicted time (from the Einstein equation (∆ *x*^2^ = 4*D* ∆ *t*)) divided by 1 μs. The 1 μs time step was chosen such that the probability of acceptance was always less than 1. Translations and rotations were considered independent.

### Graphical analysis of antennae size and PSII connectivity

The PSII-LHCII chlorophyll antenna network was modelled as a graph network using the NetworkX package in Python (Supplementary Fig. 4). Each PSII or LHCII complex was represented by a node in the graph. Edges were introduced between any two nodes in the graph if any chlorophyll in the particle represented by the first node was within a given distance (referred to as the distance threshold) of any chlorophyll in the complex represented by the other node (Supplementary Fig. 4). We measured two properties of the graph, termed the antennae size and the PSII connectivity. The antenna size was defined as the average number of LHCII, over all PSII complexes and at a given distance threshold, for which there exists a connecting path (a sequence of edges connecting the two nodes). The PSII connectivity was defined in terms of PSII clusters. If there existed a path between any two PSII complexes, they were defined as belonging to the same cluster of PSII complexes. The PSII connectivity was therefore defined as the average number of reaction centres per PSII cluster.

### Code Availability

Source code is freely available under a creative commons license at https://github.com/WhjWood/Thylakoid_Model.

## Appendix 1. A lattice-based algorithm for calculating spatial overlap of particles of arbitrary geometry

The particles were regarded as “hard” objects, meaning that translations and rotations with resulted in any two particles occupying the same lattice site were forbidden. To calculate whether particle overlap had occurred during the movement of a given particle, we developed the following algorithm (Supplementary Fig. 2)

Particle geometries, as determined from PDB structures, were represented by a matrix P_i_ of dimension 2 x n. The columns of P_i_ contain the coordinates of lattice sites occupied by the particle.

We define the matrix M as the concatenation of all *P*_*i*_ along the column axis, and is therefore a matrix which contains, in its columns, the lattice sites occupied by all (m) particles

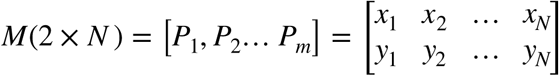

During a given time step, the i^th^ particle may be translated by an amount an amount ∆ *X* and rotated by an amount ∆ *θ*, giving the new coordinates.

The new M is created after each particle translation or rotation.

### Theorem 1.

We define the function Ν_unique_(M) as the number of unique columns of M.

*P*_*i*_ spatially overlaps at least one other particle in M if and only if

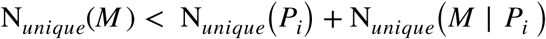

Where A|B (A but not B) refers to the set difference of the columns of A and B. Intuitively, if the total number of occupied lattice sites is less than the number occupied by the particle of interest plus the number occupied by all other particles (Figure 1A,B), then the particle of interest must share a lattice site with another particle. Therefore, there is overlap of two or more particles.

Proof.

As N_unique_(M) is equivalent to the number of elements in the set of columns of M, or N_*unique*_(M) = |{*Col*(*M*)}|, and because M contains the columns of *P*_*i*_ then

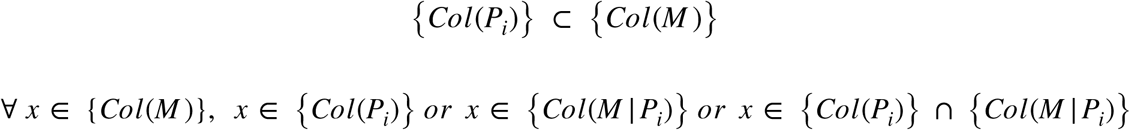

And so

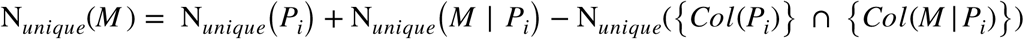

Hence

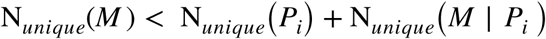

implies that

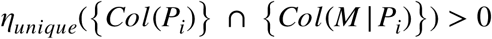

And so there exists at least one column (and therefore at least one occupied lattice site) common to both *P*_*i*_ and *M* | *P*_*i*_.

We use existing Python libraries to calculate N_unique_. We found that we could significantly reduce the time taken to compute N_unique_ if lattice sites were given a unique index, effectively making the lattice geometry 1-D. We apply suitable scaling and offsetting of the lattice units to ensure that *x*, *y* ∈ ℕ and define a suitable function f

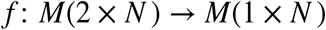

Such that the uniqueness of the columns of M is preserved. For 2 dimensions, we find a suitable f in the form

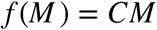

Where the 1×2 matrix C is in the form

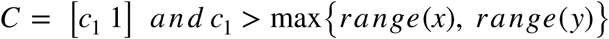

For example, if 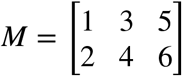 and *C* = [10 1], then *f*(*M*) = [12 34 56]. In this study, *c*_1_ = 10^4^.

### Theorem 2.

Column i of *f*(*M*) = *CM* is unique in CM if column i of M is unique in M

Proof.

We define *f*(*x*, *y*)

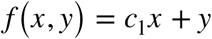

Where ∀*x*, *y c*_1_ > *x*, *y*

For *x* = 0 *f* (0,*y*) = *y*

This is unique for any y since

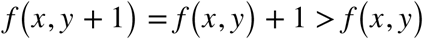

For 0 < *x*, we have *f*(*x*, *y*) = *c*_1_*x* + *y*

Now *y* < *c*_1_ and so, for all *y, f*(*x*, *y*) = *c*_1_*x* + *y* < *c*_1_*x* + *c*_1_ = *f* (*x* + 1,0)

And so

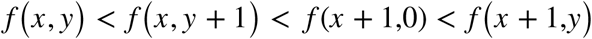

In summary, if *f* (0,0) is uniquely defined, then *f*(*x*, *y*) = *c*_1_*x* + *y* is also uniquely defined if *c*_1_ is greater than any given x or y in the geometry of the model.

Using the algorithm outlined above we were able to rapidly asses overlap (Supplementary Fig. 2). The model used in this study contained 1166 particles occupying 120600 lattice sites.

## Results

### The thylakoid model

A model of the plant thylakoid membrane was constructed, based on realistic protein complex geometries and densities (Fig. 1A). The densities of PSII and PSI were obtained from AFM images on isolated grana and stromal lamellae membranes from spinach (23). The relative number of other complexes in the membrane (LHCII, cyt*b*_*6*_*f*, ATP synthase) were taken from the current literature, see *methods*. The model produced a number of predictions that agreed with the experimental literature. The chlorophyll *a*/*b* ratio calculated to be 3.2 was found to agree with those taken from spinach leaves grown at 150 µmol photons m^−2^ s^−1^ (Supplementary Fig. 5). In this model, the grana area, PSII density, PSI density and PSI: PSII stoichiometry were fixed parameters based on experimental data. The number of PSII complexes was calculated by multiplying the PSII density by the grana area. Because of the fixed PSI:PSII ratio, this resulted in a fixed number of PSI complexes. The radius and therefore the area of the stromal lamellae was calculated such that it resulted in the correct PSI density. In this model we choose a 1:1 stoichiometry of PSI and PSII. This resulted in the radius of the thylakoid including stromal lamellae to be 252 nm when the grana radius was set at 190 nm (23) and predicted the area of grana as a percentage of total thylakoid area to be 54%. We found this prediction to be within the range of 50-80 % proposed in the literature (59, 60). In general, we found that for any choice of PSI to PSII stoichiometry in the range of 0.5 to 1.2, the resulting area of grana as a percentage of total thylakoid was within the observed range (Supplementary Fig. 6). This largely agrees with the range of PSI:PSII ratios of 0.54 - 1.4 observed in the literature (61). We calculated the area occupation of each type of complex in the membrane as the number of lattice sites occupied by the complexes divided by the total number of lattice sites in the grana or stromal lamellae. The area fraction (the fraction of lattice sites occupied) of protein complexes were 0.75 in the grana (Fig. 1B) and 0.67 in the stromal lamellae (Fig. 1C). The lower density of proteins in the stromal lamellae compared to the grana was confirmed with AFM analysis of grana and stromal lamellae membranes (Supplementary Fig. 7) and previous work (62). The overall protein area occupation as a percentage within the thylakoid was 72%, in agreement with the previous estimates of around 70 % (63). In particular, LHCII dominates the grana with free LHCII trimers accounting for almost 40 % of the grana area (Fig. 1C). Including 4 LHCII timers bound to each C_2_S_2_M_2_ PSII complex and 2 LHCII trimers bound to each C_2_S_2_ PSII complex, increases the area occupation of LHCII to around 55 % of the grana area. The area of the stromal lamellae was found to be predominantly occupied by PSI (Fig. 1D). At 40 %, the area occupancy of PSI in the stromal lamellae was greater than that other every other protein complex in the stromal lamellae combined, which was approximately 30 %.

### State transitions change the molecular organization of the thylakoid membrane

We investigated the phenomenon of state transitions at an individual complex level. In the dephosphorylated State I (SI), LHCII trimers interact with one another both within the same membrane via specific protein-protein interactions (hereafter *lateral* interactions) and across the stromal gap of the grana via electrostatic interactions mediated by cation screening (hereafter *stacking* interactions) (Fig. 2A). Both of these interactions are greatly diminished when LHCII is phosphorylated in State II (SII) (13, 14). There is however, the appearance of the PSI-LHCI-LHCII complex when LHCII is phosphorylated (Fig. 2B) (35). We modelled SI using the lateral and stacking LHCII interactions described in Schneider & Geissler (16)(Fig. 2A,C) and modelled SII by introducing a square well interaction between LHCII and PSI (Fig. 2B,D). In both cases, only LHCII was allowed to move between the grana and stromal lamellae, consistent with the majority of the biochemical data (31, 40, 42). The grana radius was set to 190 nm and 170 nm for SI and SII respectively as observed in SIM images of spinach chloroplasts (23). The densities of PSII and PSI were unchanged from SI to SII as was the area of grana as a percentage of total thylakoid area (31, 56). The latter fact is explained by the increased number of grana per chloroplast that offsets the decrease in grana diameter in State II (31). Monte Carlo simulations were used to sample the partitioning of LHCII into the grana and stromal lamellae at equilibrium. Sampling was carried out after 10^7^ Monte Carlo perturbations for SI and SII (Supplementary Fig. 1).

**Figure 2.**
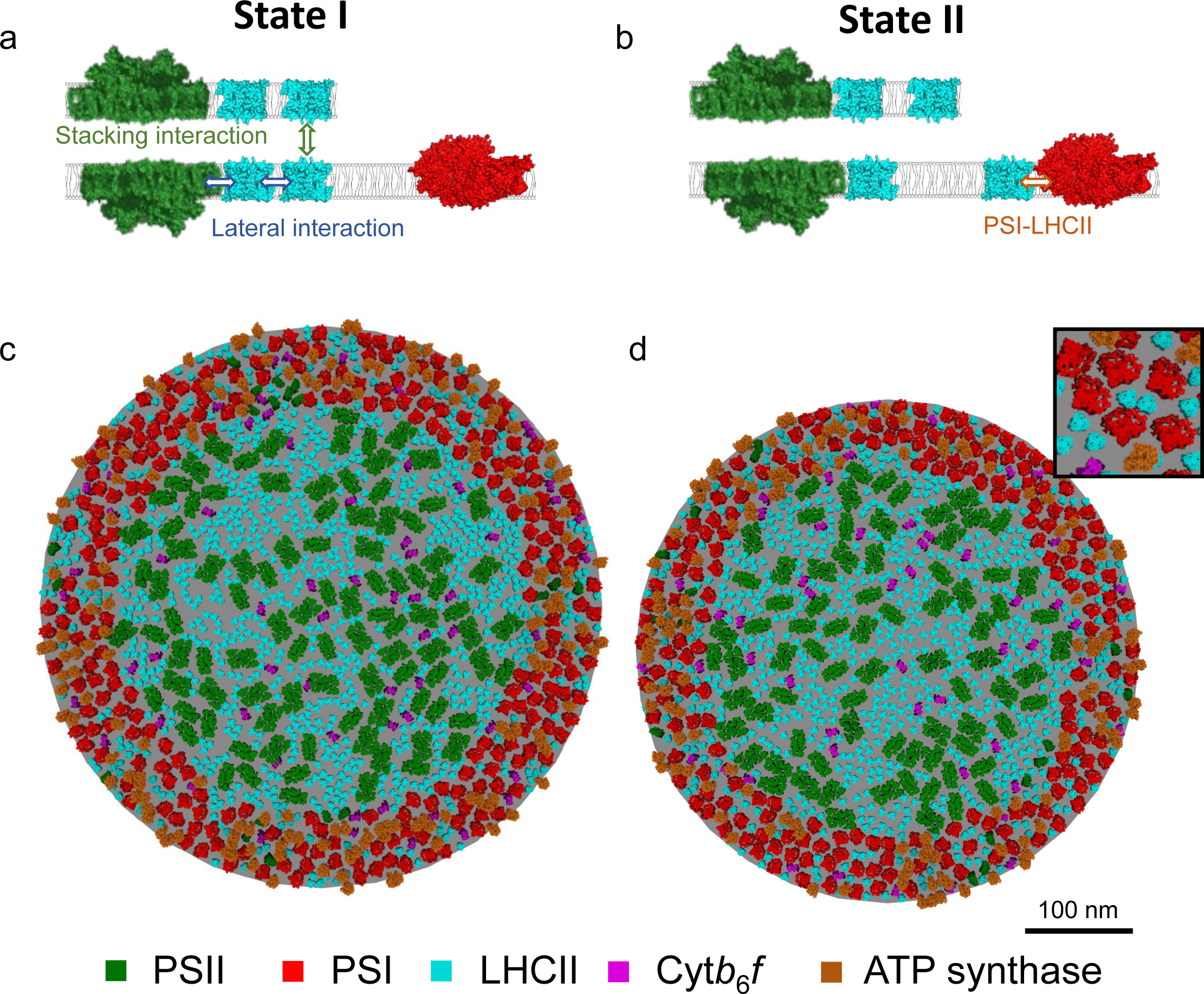
Modelling state transitions. **a**, SI is mediated by grana stacking attractive interactions and LHCII-LHCII association, including PSII-bound LHCII trimers. **b**, SII is mediated by PSI-LHCII attractive interactions only. Visual representations of the model in SI (**c**) and SII (**d**) show differences in the organization of thylakoid membrane.

We defined the presence of the PSI-LHCII complex by the criterion of LHCII in close proximity (≤ 1 nm) to the PSI binding site described in Pan *et al*., (36). We observed an increased in the fraction of PSI in complex with LHCII from 2.3 ± 0.6% at 0 K_B_T to 80% at 16 K_B_T in SII, within this range of values an interaction of 3.3 K_B_T reproduced a figure of 42 % of PSI in complex with LHCII in SII in line with experimental data (37, 64)(Fig. 3A). We therefore chose 3.3 K_B_T as representative of the PSI-LHCII interaction in SII and hereafter we will simply refer to this as SII. We found that the strength of the PSI-LHCII interaction did not affect the amount percentage of LHCII partitioned into the stromal lamellae (Fig. 3B) but transition from SI to SII was accompanied by an increase in proportion of LHCII partitioned into the stromal lamellae by 27.5% to 29.0% (Fig. 3C). We found that the migration of LHCII into the stromal lamellae did not directly result in an increased proportion of PSI-LHCII complexes. This was evident due to the random state (no interactions) having similar amount of LHCII in the stromal lamellae (Fig. 3C) but a much-decreased proportion of PSI bound to LHCII (Fig. 3D). This result suggests that the amount of LHCII present in the stromal lamellae is predominantly controlled by the strength of the lateral and stacking interactions present in SI rather than the PSI-LHCII interaction, a finding which is consistent with experimental data that shows the reduction in grana diameter and stacking in SII is not dependent on specific PSI-LHCII interactions (31). This observation also suggested that the formation of PSI-LHCII complexes in SII was not limited by LHCII migration from the grana. To test this, we ran a separate SII simulation where LHCII was not permitted to migrate between the grana and the stromal lamellae and found that the percentage of PSI present as PSI-LHCII complexes was not significantly reduced (Fig. 3E). Thus, the partitioning of LHCII between the grana and stromal lamellae did not account for the magnitude of the increase in the number of PSI-LHCII supercomplexes as a fraction of the total number of PSI complexes in SII. It is possible that the migration of LHCII in the stromal lamellae is restricted due to the high density of the stromal lamellae. To test whether the number of PSI-LHCII complexes would increase in SII if the density of proteins in the stromal lamellae was lower, we increased the area of the stromal lamellae, effectively reducing the density by half, whilst keeping the grana area the same (Fig. 3F). This led to a large increase in the percentage of LHCII in the stromal lamellae, reaching 42% in SII. Counterintuitively, the decreased density also led to a decrease in the number of PSI-LHCII complexes from 42 % to 6.4 % (Fig. 3F). We speculate that this decrease may be due to an increased entropic cost of forming the PSI-LHCII complex when the density of the stromal lamellae is low.

**Figure 3.**
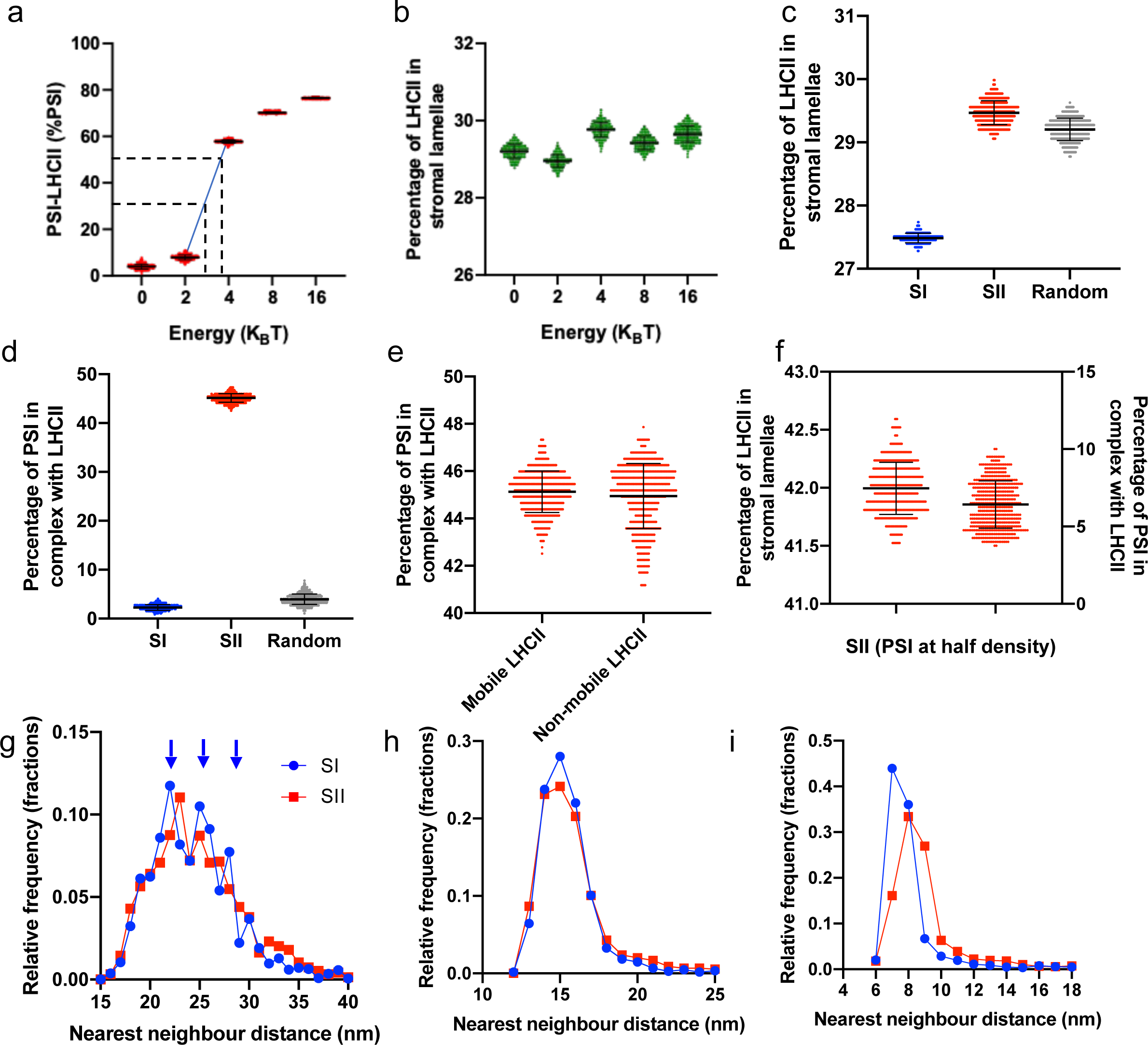
LHCII interactions control the partitioning of LHCII between the grana and the stromal lamellae and affect the organization of PSII but not PSI. **a**, The percentage of PSI in complex with LHCII with varying strength of the PSI-LHCII interaction (dashed lines show experimentally observed range of %PSI-LHCII). **b**, The percentage of LHCII in the stromal lamellae with varying strength of the PSI-LHCII interaction. **c**, The percentage of LHCII complexes partitioned into the stromal lamellae in SI (blue), SII (red) and with no interactions present (random, grey). **d**, Percentage of PSI in complex with LHCII (defined as LHCII ≤ 1 nm from binding site) in SI, SII and random simulations. **e**, Percentage of PSI in complex with LHCII in SII when LHCII is or is not allowed to migrate between the grana and the stromal lamellae. **f**, The percentage of LHCII complexes partitioned into the stromal lamellae and the percentage of PSI in complex with LHCII after the density of the PSI in the stromal lamellae was decreased by half (this was achieved by increasing the area of the stromal lamellae). **g**, The nearest-neighbour distribution (centre-to-centre distances) of PSII-LHCII supercomplexes in SI (blue circles) and SII (red squares). The 3 modes arising in SI are indicated by blue arrows. **h**, The nearest-neighbour distribution (centre-to-centre distances) of PSI in SI and SII. **i**, The nearest-neighbour distribution (centre-to-centre distances) of LHCII in SI and SII.

We next investigated whether the movement of LHCII led to reorganization of the thylakoid membrane, by analysing the nearest-neighbour distributions of PSII (including C_2_S_2_M_2_ and C_2_S_2_ supercomplexes). In our model PSII-LHCII did not form grana-spanning semi-crystalline domains, as has been seen previously (52), however we did observe a more ordered arrangement of PSII supercomplexes in SI (Fig. 3G). The trimodal nearest-neighbour distribution of PSII in SI (Fig. 3G, blue arrows) is in agreement with the organization of PSII observed in AFM images of grana membranes in SI (7, 43, 65) as is the more disordered arrangement in SII (23, 56). Despite the increase in the number of LHCII and PSI-LHCII complexes in the stromal lamellae in SII compared to SI, PSI nearest-neighbour distributions were indistinguishable (Fig. 3H). This is in contrast with the observed increased amount PSI dimerization is SI observed in AFM images of stromal lamellae membranes (23). Here, as with PSII, it is difficult to ascertain at present as to whether the discrepancy between the results of the model with previous experiments is due to artefacts resulting from the detergent-based isolation of membranes in Wood et al (23) or that the dimerization of PSI is due to interaction not included in the model. We note that Wietrzynski et al., (8) using cryo ET found no evidence of such PSI dimers in *Chlamydomonas* thylakoids though the PSI structure is different to plants (66). A large difference between SI and SII was observed in the nearest neighbour distribution of LHCII (Fig. 3I). The mean nearest neighbour distances of LHCII in SI and SII were found to be 8.27 nm and 9.02 nm respectively. This can be seen in the clustering of LHCII in SI (Fig. 2C) compared to the seemingly more random LHCII organization in SII (Fig. 2D).

### A diffusion-based lower bound for the time taken for the transition from state I to state II

The transition from SI to SII relies on the phosphorylation of LHCII (26). This is followed by the diffusion of a portion of LHCII into the stromal lamellae. However, which of the two processes, phosphorylation or diffusion is the slowest and therefore rate-limiting is unclear. We performed time-resolved simulations with the intention of estimating the time taken for a state transition to occur if LHCII phosphorylation was immediate (the diffusion limited case). We could not simulate the directly the SI to SII transition in a time resolved manner as this would require a kinetic Monte Carlo simulation which is computationally unfeasible for this model. Instead, we substituted a model which lacked any interactions (hereafter referred to as a random distribution) in place of SII, which, in relation to the amount of LHCII in the stromal lamellae, is highly similar to SII (Fig. 3C). The diffusion coefficients of the thylakoid protein complexes were calculated using the Saffman-Delbrück model (57)(see methods). Table 1 shows the calculated translational (D_T_) and rotational (D_R_) diffusion coefficients used in the simulations. We found the half-time for the transition from SI to random as measured by the proportion of LHCII in the stromal lamellae was 0.95 ms (Fig. 4A). This is much faster than state transitions occur *in vivo*, which have a half-time of around 3-8 minutes (34), suggesting the limiting factor in the time taken for state transitions to occur is the action of the STN7 kinase. By analysis of the mean square displacement of LHCII in the random state, we calculated the effective diffusion coefficient of LHCII to be 1.6 × 10^−10^ cm^2^ s^−1^ (Fig. 4B). This is within the range observed for LHCII and phosphorylated LHCII *in vivo* which were found to be 8.4 × 10^−11^ cm^2^ s^−1^ and 2.7 × 10^−10^ cm^2^ s^−1^ respectively (67). It is evident in this model that LHCII migration into the stromal lamellae is driven by the higher density of protein complexes in the grana in SI. During the course of the transition, LHCII continues to flow from the grana into the stromal lamellae until they are almost at equal area densities (Fig. 4C).

**Table 1.** Effective radii, translation diffusion coefficients D_┬_ and rotational diffusion coefficients D_r_ calculated using the Saffman & Delbruck model (1975)

| Particle | Effective radius (nm) | $D_T$ ($\times 10^{-9} \text{ cm}^2 \text{ s}^{-1}$) | $D_r$ ($\times 10^3 \text{ rad}^2 \text{ s}^{-1}$) |
| --- | --- | --- | --- |
| PSII (C2S2M2) | 12.7 | 1.46 | 0.25 |
| PSII (C2S2) | 10.1 | 1.55 | 0.40 |
| B6F | 5.3 | 1.82 | 1.50 |
| LHCII | 4.5 | 1.88 | 2.00 |
| PSI | 7.2 | 1.69 | 0.78 |
| ATP Synthase | 3.7 | 1.97 | 3.06 |
| PSII (monomeric) | 5.8 | 1.78 | 1.20 |

**Figure 4.**
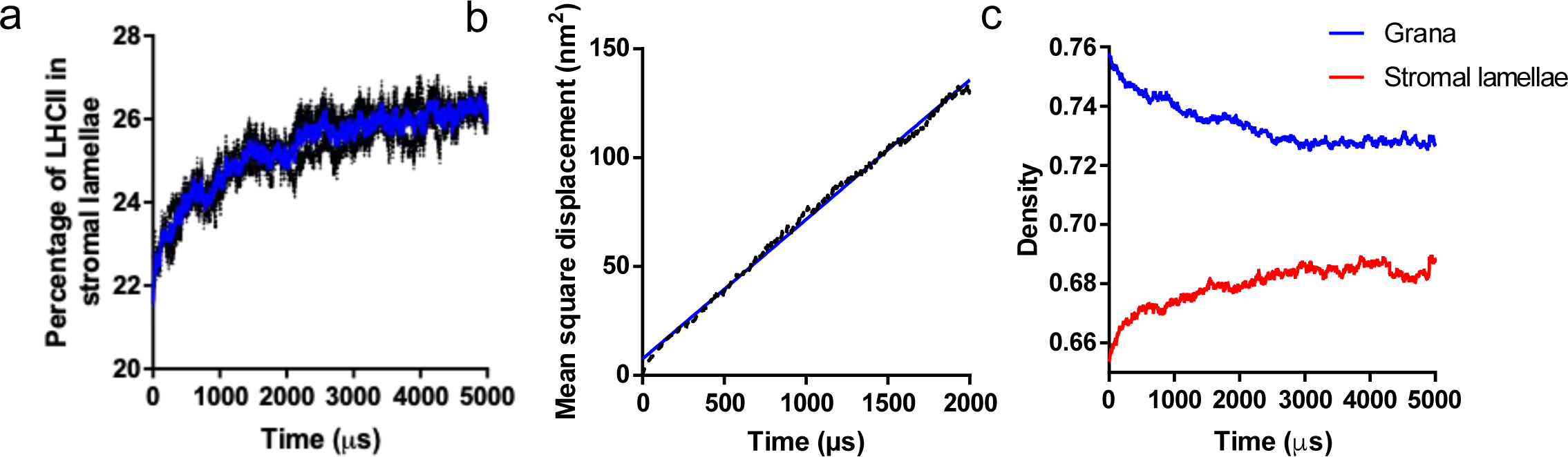
The diffusion-based timescale for the migration LHCII into the stromal lamellae is sub-millisecond. **a**, Kinetics of the percentage of LHCII in the stromal lamellae during the transition from SI to the random state (blue solid line: average of 3 traces; black dashed lines: individual traces). **b**, The mean square displacement of LHCII over time (blue solid line: linear fit; black dashed line: raw values). **c**, The kinetics of the density of the grana (blue) and stromal lamellae (red) during the transition from SI to the random state.

### Lateral and stacking interactions of LHCII cooperate to increase the light-harvesting antenna size and the connectivity of PSII-LHCII in SI

We investigated the effects of the lateral and stacking interactions of LHCII individually, to ascertain their impacts on the properties of the thylakoid membrane. Simulations were carried out containing either both lateral and stacking interactions, lateral interactions only, stacking interactions only, or no interactions. We found that the lateral interactions caused the LHCII complexes to cluster as shown by a decrease in the nearest neighbour distances (Fig. 5A). Stacking interactions did not cause clustering of LHCII complexes in the same layer (Fig. 5A) but the attraction between LHCII complexes on opposing layers did lead to increased correlation between LHCII complexes as shown by a decreased nearest neighbour distance when considering only neighbours on the opposing thylakoid layer (Fig. 5B). These results were also replicated in simulations containing LHCII only (Supplementary Fig. 3).

**Figure 5.**
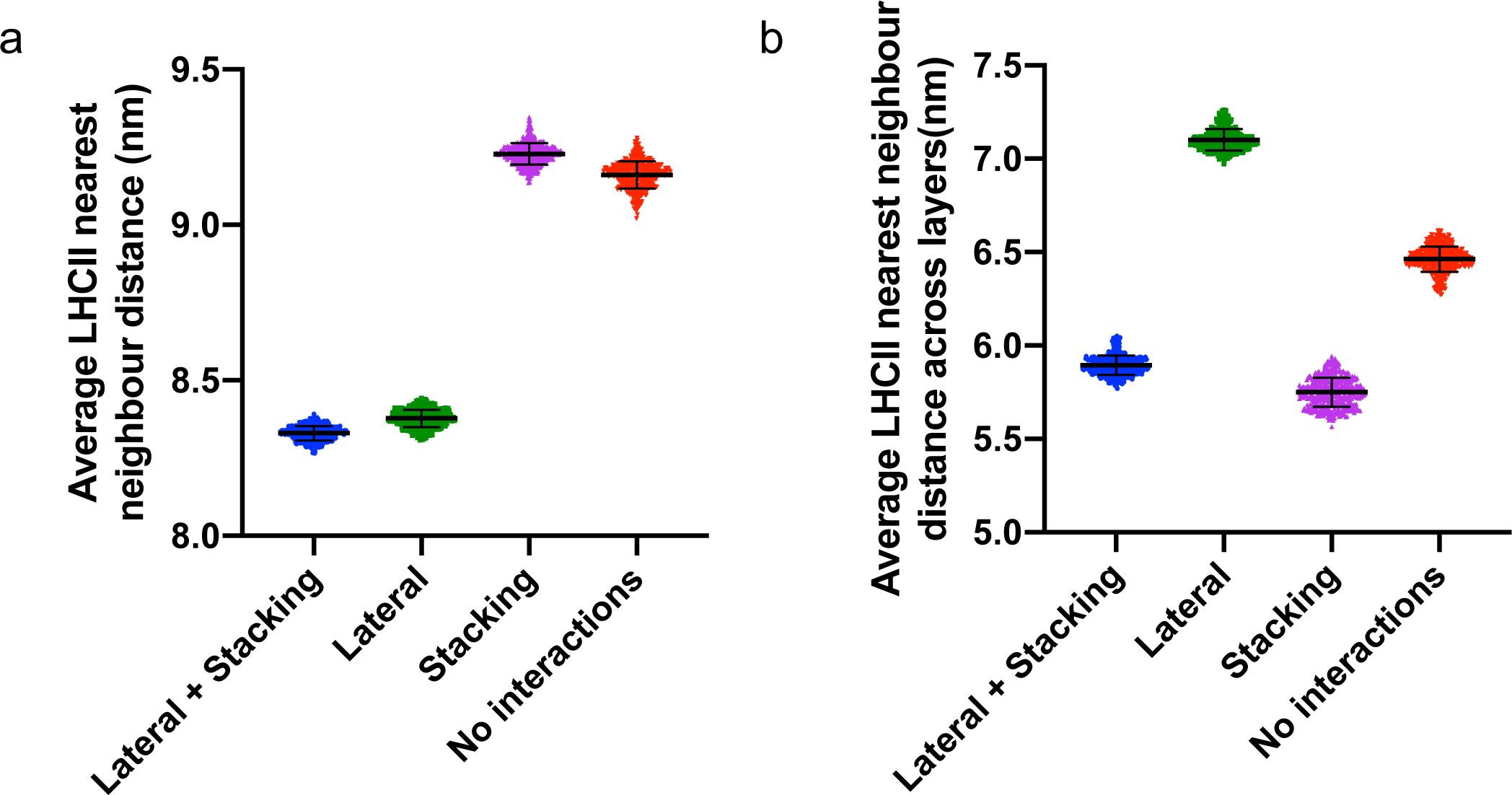
Lateral and stacking interactions affect the organization of LHCII. **a**, Average nearest neighbour distances of LHCII with the same layer in simulations containing lateral and stacking interactions (blue), lateral interactions only (green), stacking interactions only (magenta), or no interactions (red). **b**, Average nearest neighbour distances between LHCII complexes in layer 1 and layer 2 (across layers) in simulations as described in (a).

Questions naturally arise from the simulations outlined above as to how the increased density and clustering of LHCII in the grana in SI are implicated in the observed increase of antenna size and connectivity of PSII observed *in vivo* (33, 68). We investigated the structure of the PSII-LHCII chlorophyll light-harvesting antenna using a network-based topological analysis. The antenna network consisted of LHCII trimers connected to PSII reaction centres (Fig. 6), which collectively made up the vertices of the graph. Edges between neighbouring LHCII trimers and/or PSII supercomplexes were introduced if there were chlorophylls in each complex close enough together. That is, the Euclidean distance was less than a given value called the distance threshold (Supplementary Fig. 4, Fig 6A). Reaction centres where assumed to be connected to bound LHCII trimers within the same C_2_S_2_M_2_ or C_2_S_2_ PSII-LHCII supercomplex. The quantitative properties of the antenna network are highly sensitive to the chosen distance threshold for this type analysis. For this reason, we describe the phenomena observed for a range of distance thresholds and show that the qualitative conclusions drawn are independent of the distance threshold. Firstly, for any given distance threshold, we found that PSII and LHCII network was arranged in functional clusters, with a defined number of reaction centres and associated LHCII trimers akin to the heterogeneous photosynthetic unit model of Lavergne & Trissl (69). At low distance thresholds (≤ 1 *nm*), one retrieves the average antenna properties of the C_2_M_2_S_2_ and C_2_S_2_ supercomplexes (at equal stoichiometry), namely around three LHCII trimers and two reaction centres per cluster. With lateral and stacking interactions together or only lateral interactions, the size of the clusters was increased, leading to a larger antenna size (Fig. 6B) and more PSIIs per cluster (Fig. 6C). The increase in number of reaction centres per cluster is in agreement with the decrease in PSII connectivity observed in SI by Kyle *et al* (33). We found only a slight increase in the antennae size and the number of PSIIs per cluster when only stacking interactions were present (Fig. 6B, C) compared to when no interactions were present. Using the same approach, we investigated the structure of the PSI-LHCII chlorophyll network (Fig. 6D). We observed a slight increase in the network-based PSI antenna size (Fig. 6D) in SII and no interactions compared to SI. This can be seen to occur at low (< 3 nm) and high (> 5 nm) distance thresholds but not in the intermediate range.

**Figure 6.**
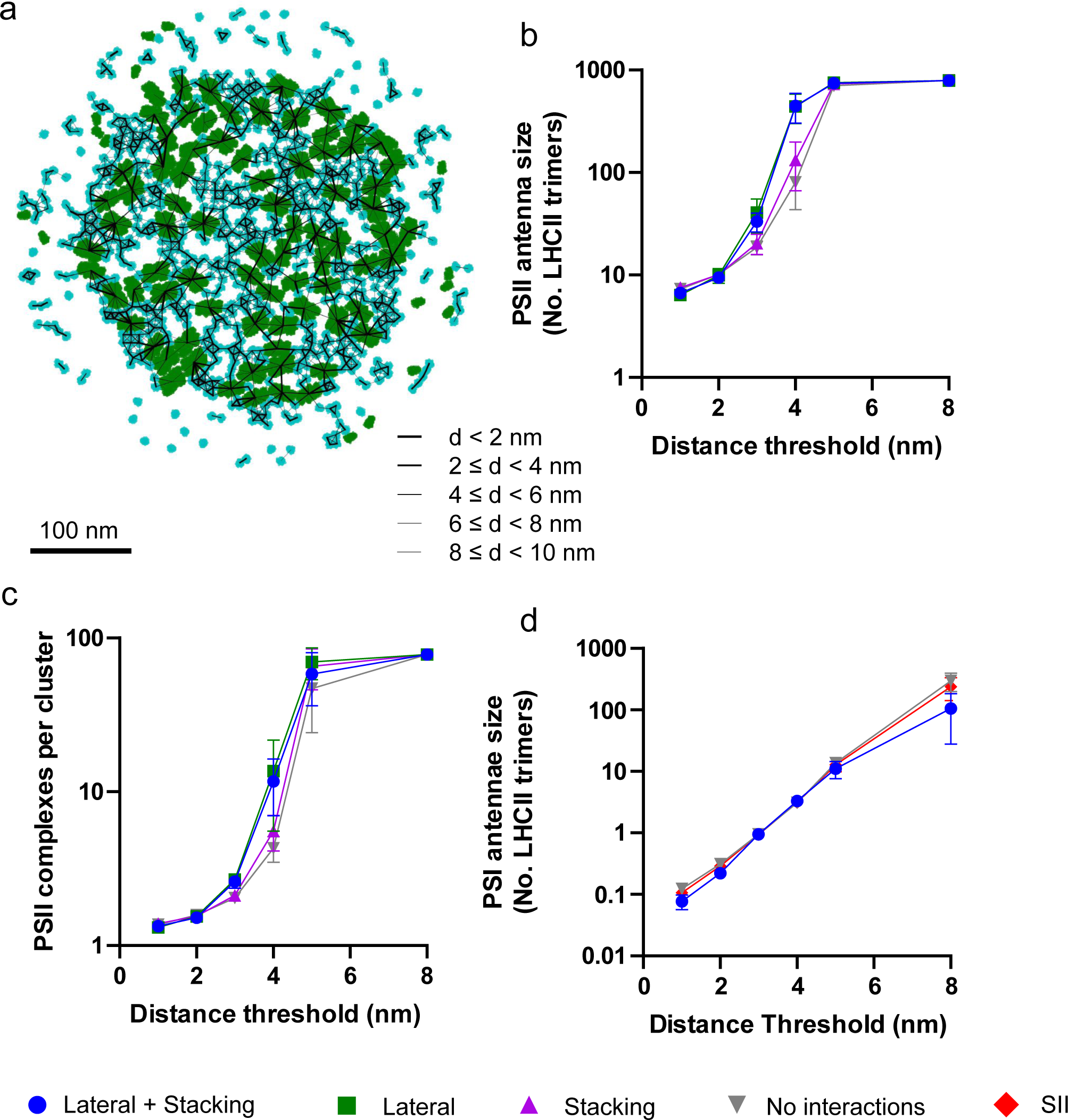
Analysis of changes in the structure of the chlorophyll light-harvesting antennae network resulting from lateral and stacking interactions. **a**, Chlorophyll network produced by simulations containing lateral and stacking interactions. PSII and LHCII complexes are shown in green and cyan respectively. Black lines between the centres of any two complexes indicate that there exists a chlorophyll on one complex less than 10 nm from a chlorophyll on the other and are weighted by the distance (shorter distances (< 2 nm) are shown by the thickest lines). **b**, Number of LHCII complexes per PSII in the presence of lateral and stacking interactions (blue circles), lateral interactions only (green squares), stacking interactions only (magenta triangles), or no interactions (inverted grey triangles) **c**, Number of PSII complexes per cluster resulting from simulations labelled in accordance with (b). **d**, Number of LHCII complexes per PSI is SI (blue circles), SII (red diamonds) and no interactions (inverted grey triangles).

## Discussion

Recent reports suggest a dual role of STN7-dependent LHCII phosphorylation. On the one hand, LHCII phosphorylation leads to structural changes in the thylakoid membrane in which grana diameter and the number of membrane layers per stack decreases, while the number of grana per chloroplast increases (23, 31, 44). This process is driven by a increase in the repulsive electrostatic force between LHCII trimers across the stromal gap of appressed grana membranes (14). On the other hand, LHCII phosphorylation has been implicated in balancing the excitation energy distribution between PSII and PSI as a consequence of state transitions (21, 26). In addition to STN7-dependent LHCII phosphorylation, state transitions require specific interactions between LHCII-P and the PsaL/H/O subunits of PSI, which lead to the formation of the PSI-LHCII supercomplex (34–36). In this study we developed a molecular model of the thylakoid membrane as a basis for Monte Carlo simulations to investigate the role of lateral intermolecular interactions, changes in grana stacking forces and specific PSI-LHCII interactions in modifying the distribution of photosynthetic complexes between the two domains and the relative antenna sizes of PSI and PSII during state transitions. The value of a theoretical model is the ability to clearly separate the effects of these three forces and understand the extent to which they likely cooperate to bring about the changes observed *in vivo*. Our results show three forces act largely separately to bring about specific facets of the structural changes observed up state transitions.

The first question we addressed was how the three forces affected the distribution of LHCII between the grana and stromal lamellae. In the absence of the lateral and stacking interactions percentage of LHCII in the stromal lamellae rises due to the lower density of complexes in this region. The increase we observe explains the tendency for a slight increase in the chlorophyll *a/b* ratio of the stromal lamellae (40). The strength of the PSI-LHCII interactions however did not cause appreciable changes in the percentage of LHCII in the stromal lamellae. This finding is in line with experimental results that show that while mutants lacking the PsaL/O/H LHCII-P binding site on PSI lack state transitions they still show a transition to smaller grana in SII indicative of a movement of LHCII from grana to stromal lamellae (31). The next question was how the number of PSI-LHCII supercomplexes formed was affected. As the specific PSI-LHCII interaction was increased from 0 to 16 K_B_T the number of PSI-LHCII supercomplexes increased to from 0 to 80%. *In vivo* 30 - 50 % of PSI are found in PSI-LHCII supercomplexes (37, 42, 56). We therefore chose 3.3 K_B_T, which resulted in 42 % of PSI in complex with LHCII, as representative of SII in our simulations. Interestingly, the number of PSI-LHCII supercomplexes was not significantly higher even if the protein density of the stromal lamellae was reduced by 50%, which increased the LHCII percentage in the stromal lamellae to 42%. The availability of LHCII in the stromal lamellae does not limit the amount of PSI-LHCII supercomplexes that are formed and thus a large scale redistribution of LHCII between grana and stromal lamellae is not necessary for achieving an increase in PSI antenna size, a finding consistent with previous studies (40, 41). Indeed even in SI significant amounts of LHCII are found in the stromal lamellae (37, 42). Our model does not take into account PSI-LHCII complexes involving the weaker digitonin-sensitive LHCA binding sites (37, 38) which may further augment PSI antenna size.

If redistribution of LHCII between grana and stromal lamellae upon loss of lateral and stacking interactions is not required for the observed increase in PSI antenna size might it serve another purpose? Two other phenomena associated with LHCII phosphorylation are the decrease in connectivity between PSII reaction centres (33, 68) and the faster diffusion of the electron carriers PC and PQ/PQH_2_ between the grana and stromal lamellae (23). We found that the cooperative effects of the lateral and stacking interactions led to larger average PSII-LHCII cluster size and, moreover, more LHCII trimers per reaction centre in each cluster. This was not due to a mere increase in the LHCII in the grana, which only increased by 5 %. Instead we suggest the combined effect of these interactions is the more efficient recruitment of free LHCII trimers in the grana into the functional PSII-LHCII clusters. The decreased light harvesting capacity of PSII (antenna size and connectivity) when grana stacking interactions were removed (Fig. 6B,C) has been observed *in vivo* (33). The effect has primarily been attributed to the mixing of PSII and PSI in destacked thylakoids and the consequent spillover of energy from PSII to PSI (32, 70). Previous hypotheses as to the purpose of the evolution of grana in plants have dismissed the idea that the role of grana is to prevent excitation energy spillover (71) on the basis that cyanobacteria lack grana yet are seemingly not largely afflicted by spillover. However, we have shown that stacking interactions may increase photosynthetic efficiency even when strict separation of PSII and PSI is maintained. The findings presented here reopen this debate as they have shown that an evolutionary advantage of grana stacking could be an increased light harvesting efficiency of the PSII photosynthetic unit by more efficient recruitment of LHCII. Since this state represented that of dephosphorylated LHCII, this increased cluster size may be important in very low and high white light levels. At these extremes, the STN7 is inactivated causing dephosphorylation of LHCII (29, 30). LHCII clustering has been previously observed in high-light (72). Consequently, a large PSII antenna size allows for efficient photoprotection in high light without undermining photosynthetic capacity (68). We propose that in very low light, the increased PSII-LHCII cluster size enables PSII to capture and utilise light energy more efficiently. Understanding the contribution of smaller grana upon LHCII phosphorylation to more efficient photosynthetic electron transfer by speeding up diffusion of electron carriers between the domains requires further investigation but this maybe another synergistic benefit of LHCII phosphorylation, augmenting the effects of state transitions and PSII connectivity (23).

In conclusion our model reveals that LHCII phosphorylation, that is the exchange of LHCII-LHCII and LHCII-PSII lateral and stacking interactions for specific PSI-LHCII interactions affects the antenna structure of the two photosystems differently. While the change in PSII is an emergent property of large-scale network structure the change in PSI is modulated largely by individual LHCII binding events. This idea is consistent with the fact that PSII forms an interconnected photosynthetic unit (antenna lake) whereas PSI connectivity is much more limited (73). Several testable hypotheses emerge from our model-namely that PSII antenna connectivity and perhaps changes electron transfer kinetics brought about by weakened lateral and stacking interactions may be modulated independently of specific P-LHCII binding sites on PSI. Future experimental work is now needed to clarify the extent to which this is true.

## Author contributions

W.H.J.W. carried out the simulations. M.P.J. and W.H.J.W. co-wrote the manuscript.

## Acknowledgments

M.P.J. acknowledges funding from the Leverhulme Trust grant RPG-2016-161 and Grantham Centre for Sustainable Futures.

---

**Supplementary Figure 1.**
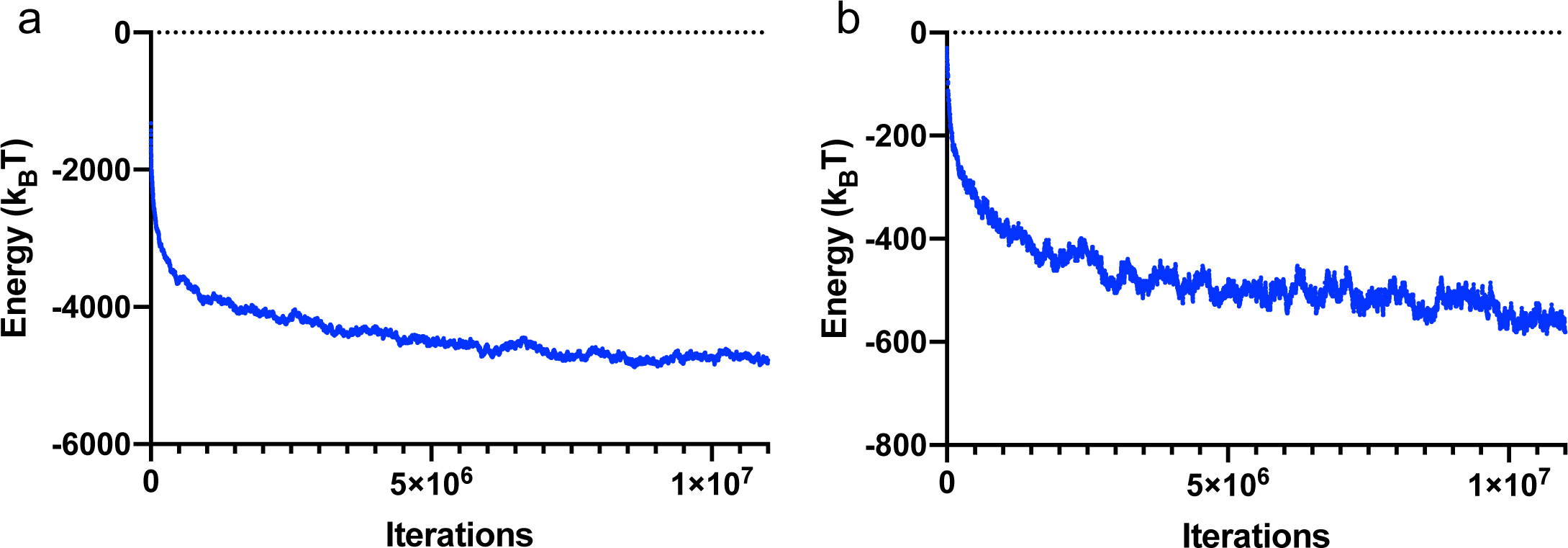
Equilibration of model prior to Monte Carlo sampling. Samples were taken after 10^7^ iterations to ensure the system had reached equilibrium. This corresponds to 4.5 and 5.6 time constants (Tau) of the energy decay with the number of iterations for SI and SII respectively.

**Supplementary Figure 2.**
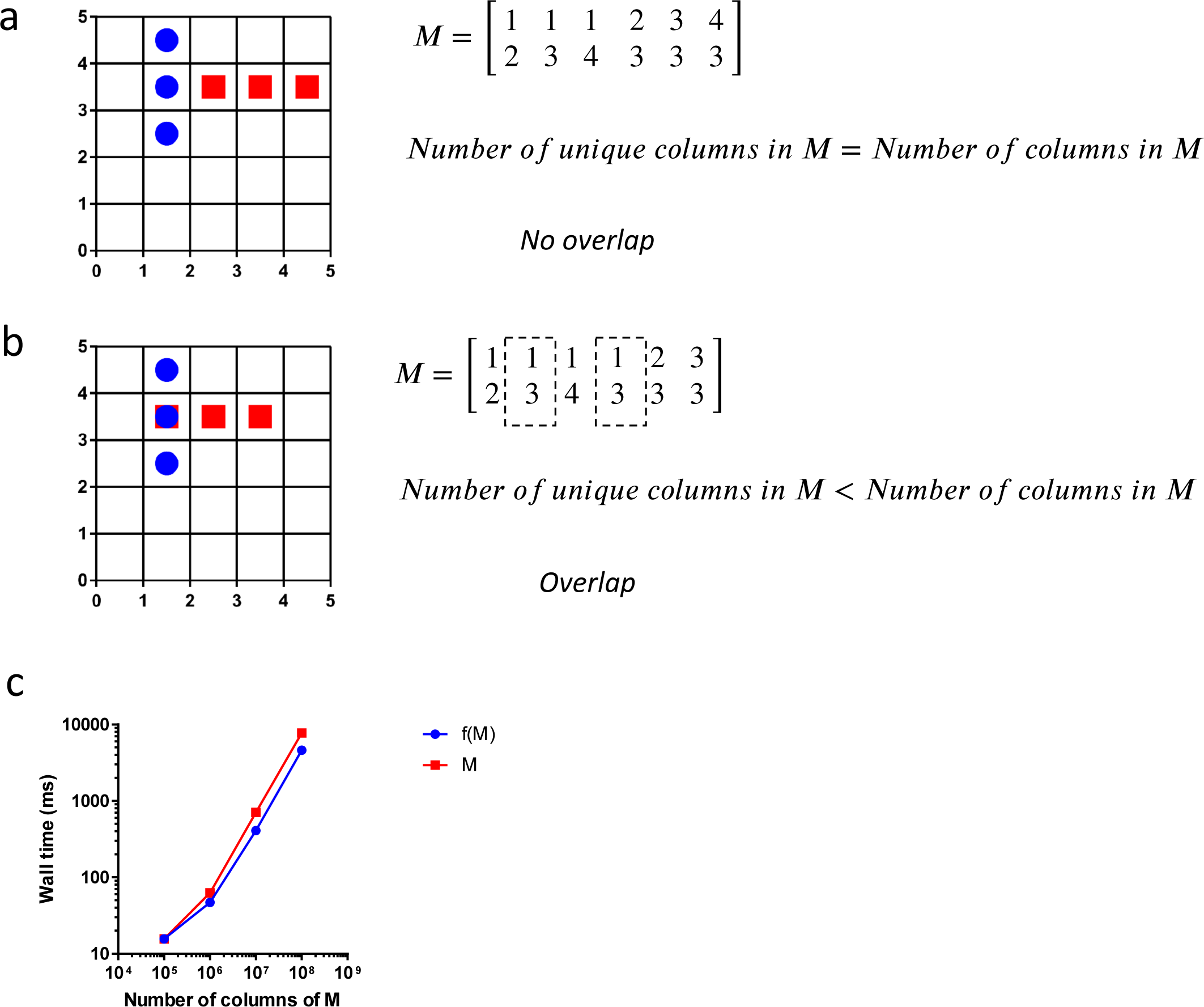
Overlap algorithm for particles of arbitrary geometry. **a**, Particle one (blue circles) does not overlap particle two (red squares). In this case, the number of unique columns in the M matrix (which contains the lattice sites occupied by all particles) is equal to the total number of columns in M. **b**, Here, particle one and particle two share a lattice site (1,3) and so they overlap. As a consequence the number of unique columns of M is less than the total number of columns of M. **c**, The scalability of the overlap algorithm applied to M (red) and the transform f(M) which transforms M into a 1-D vector (blue).

**Supplementary Figure 3.**
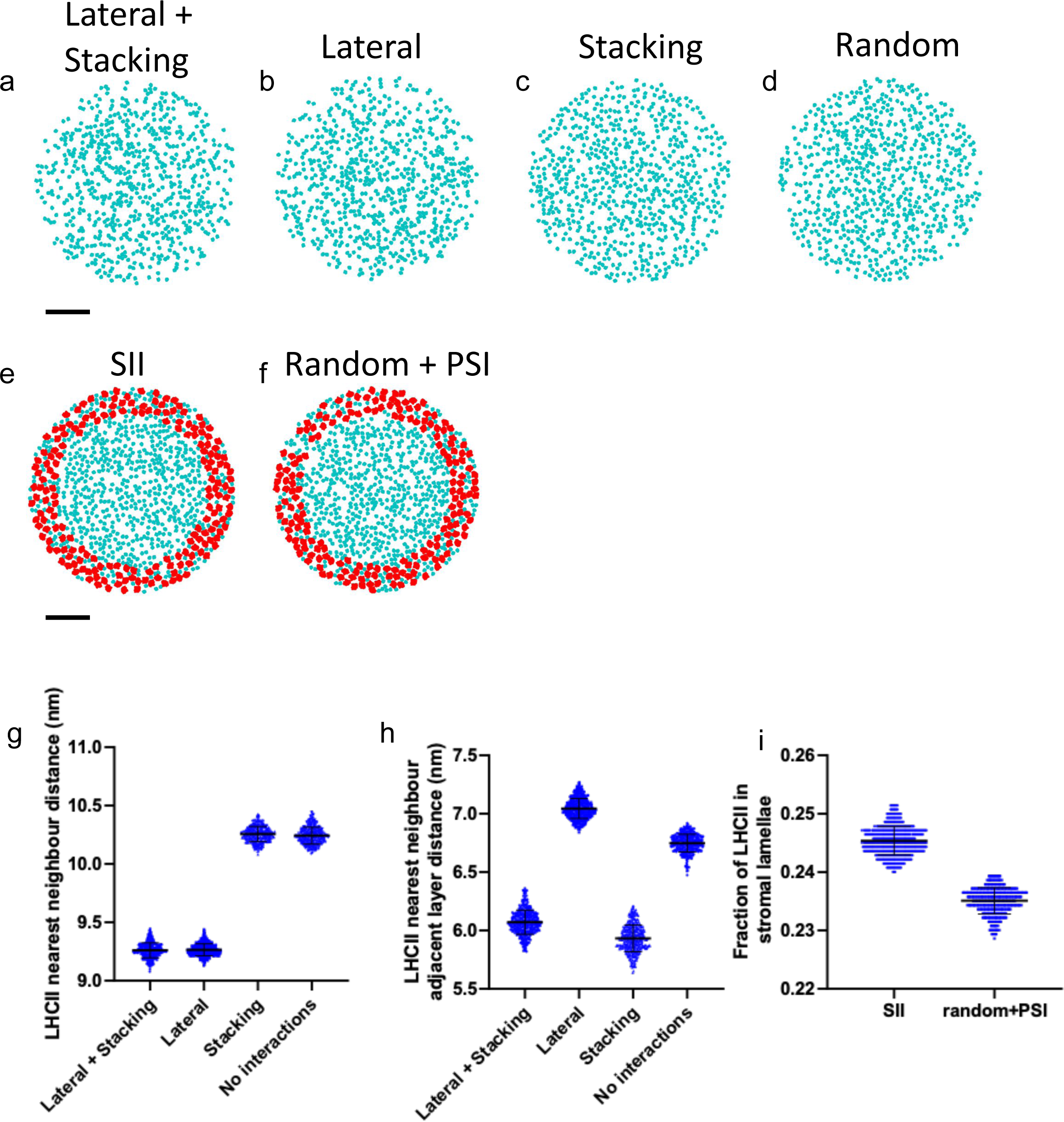
Test simulations with minimal components. **a-d**, Images from LHCII only simulations with lateral and stacking interactions (a), Lateral interactions only (b), Stacking interactions only (c), and no interactions (random, d). **e,f**, images from LHCII and PSI only simulations with PSI-LHCII interactions (SII, e) or not interactions (f). **g**, Nearest neighbour distances for LHCII particles in the same layer from simulations shown in a-d. **h**, Nearest neighbour distances for LHCII particles in adjacent layers from simulations shown in a-d. **i**, Fraction of LHCII in the stromal lamellae from simulations in e,f.

**Supplementary Figure 4.**
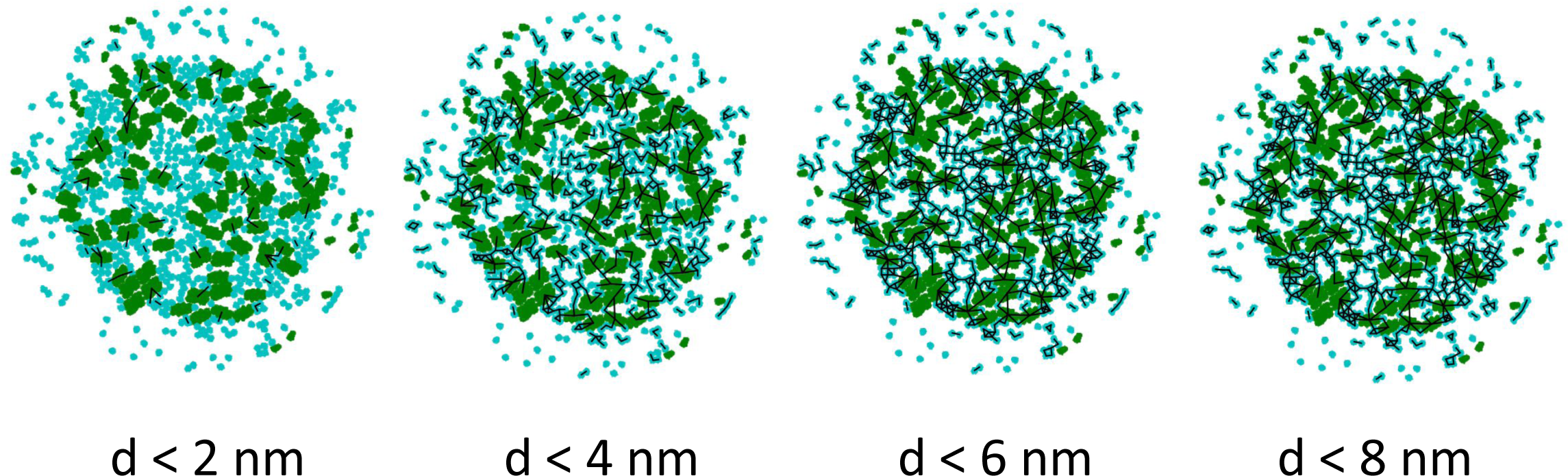
Construction of Chlorophyll networks. Edges (black lines) between any two protein complexes (LHCII and PSII) are assigned if any chlorophyll on one complex is less than a certain distance called the distance threshold (d) from any chlorophyll on the other complex. As the distance threshold is increased, the chlorophyll network becomes more connected. We investigated the structure of the networks arising from simulations containing lateral and/or stacking interactions over a range of distance thresholds (2-8 nm shown above).

**Supplementary Figure 5.**
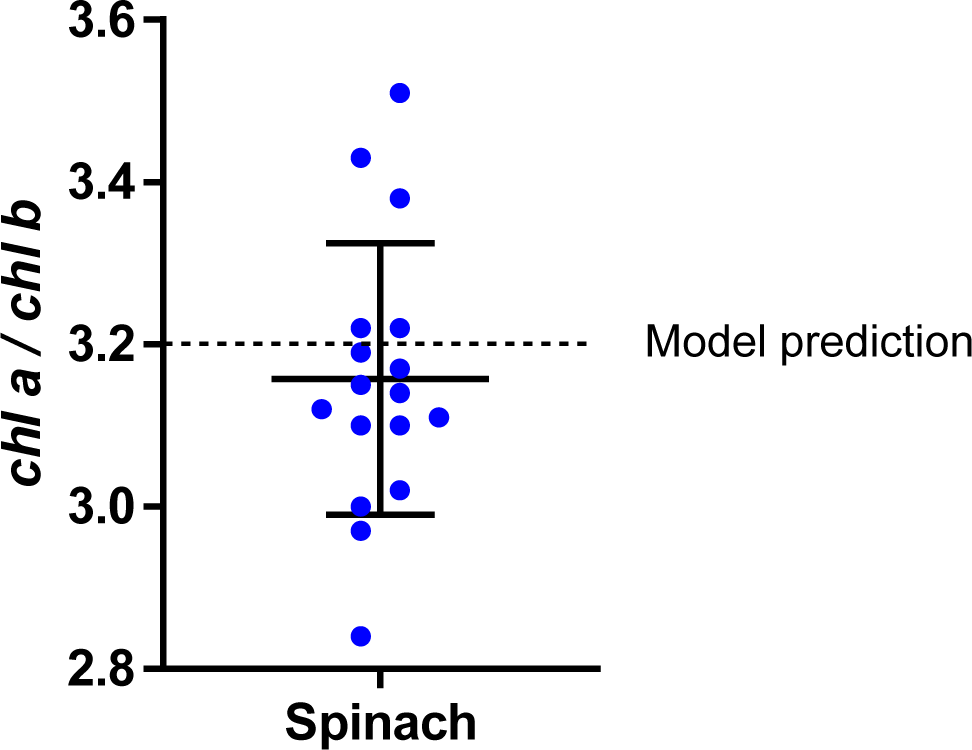
Measured ratio of chlorophyll a to chlorophyll b in Spinacia oleracea. Error bars represent mean and standard deviation.

**Supplementary Figure 6.**
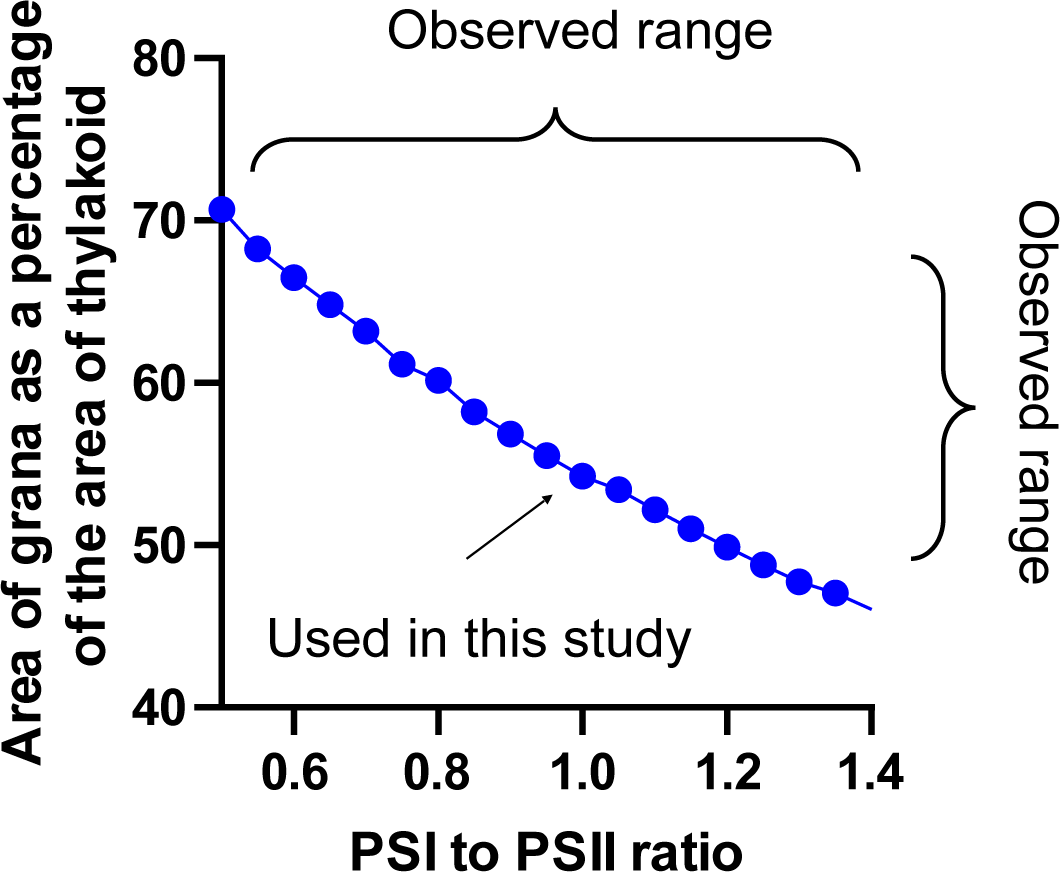
The area of grana as a percentage of total grana area is determined by the PSI to PSII ration in this model. A PSI to PSII ratio of 1 was used in this study, resulting in a thylakoid composed of 55% grana by area.

**Supplementary Figure 7.**
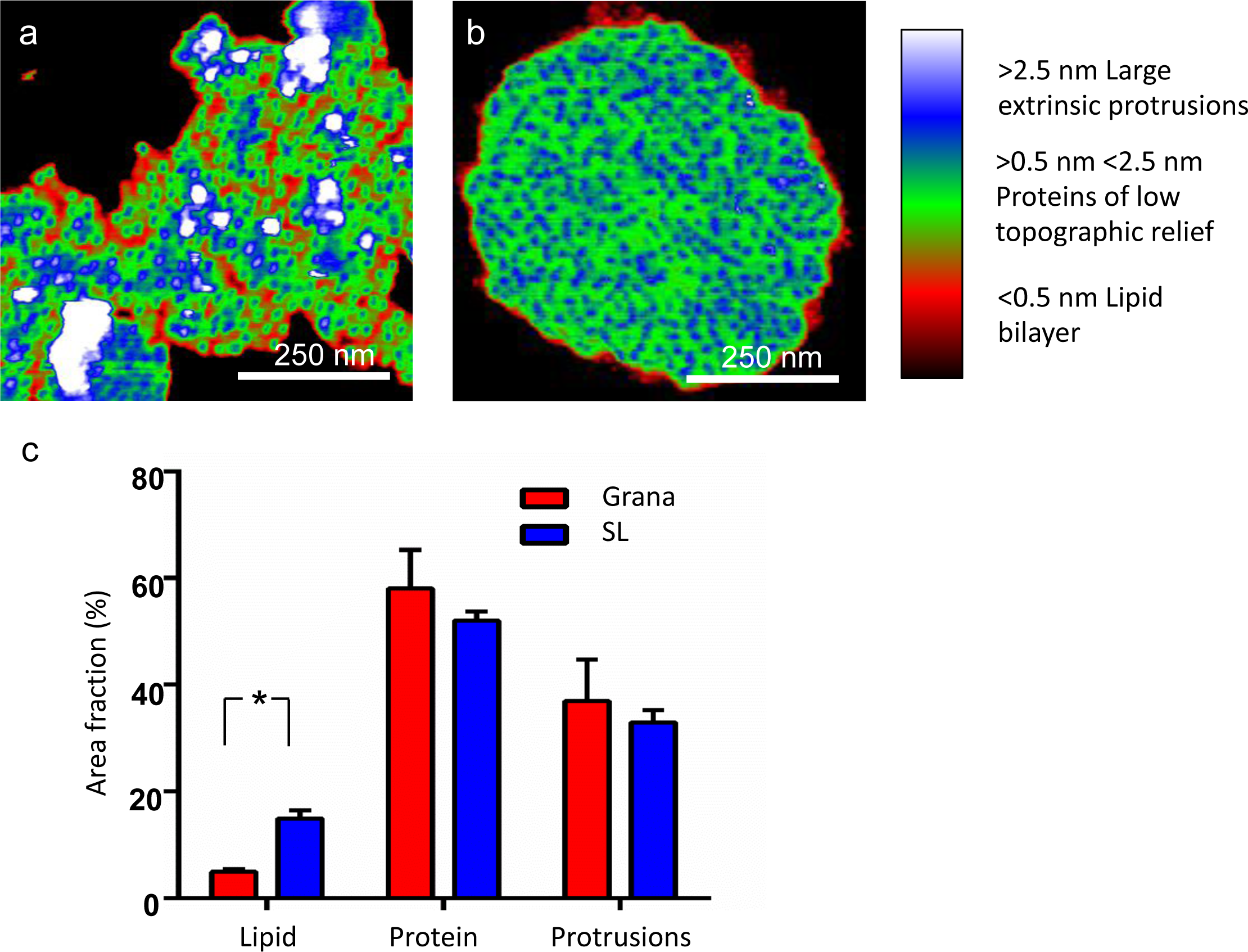
AFM analysis of the density of grana and stromal lamellae membranes. **a,b**, AFM images of stromal lamellae (A) grana (B) with colour threshold to show lipid zones in red (regions <0.5 nm above the membrane surface), proteins of low topographic relief (regions >0.5 nm and <2.5 nm above the membrane surface) and large extrinsic membrane protrusions (regions >2.5 nm above the membrane) in blue/white. **c**, A comparison of lipid, protein and protruding regions in grana (red) and stromal lamellae (blue) thylakoids (displaying mean and standard error, N = 5, * indicates p ≤ 0.001 by t-test).

